# Defining dysfunction due to loss of MECP2 in Rett Patient Brain

**DOI:** 10.1101/2021.08.24.457297

**Authors:** E Korsakova, A Morales, T McDaniel, A Lund, B Cooper, F Ma, TF Allison, K Plath, NA Graham, A Bhaduri, WE Lowry

## Abstract

Rett Syndrome is characterized by a postnatal loss of neurophysiological function and regression of childhood development. Because the syndrome is X-linked and males with MECP2 mutations generally do not survive birth, the study of this syndrome has been complicated by the fact that in female brain, a portion of neurons express wild type MECP2, and another portion express a non-functional allele of MECP2. Therefore, bulk-RNA-sequencing of Rett brain is confounded by the presence of chimerism of neurons for functional MECP2 in neurons. We developed an approach that allows for single-nuclei transcriptional profiling of individual neurons and a direct comparison between neurons that express functional MECP2 with those that express the disease-causing allele. We found that mutant neurons from Rett brain show patterns of aberrant expression of synaptic and metabolic genes, both of which can be detected in *in vitro* models of Rett Syndrome. We used these resources to identify a role for POU2F1/OCT1 transcription factor in mediating the response to stress due to loss of MECP2, highlighting a potential key molecular regulator of stress in Rett neurons. Together, our new sorting approach enables us to highlight defective molecular and metabolic pathways in Rett brain neurons and suggests that *in vitro* models could serve as valuable tools to further study this syndrome and potentially for development of novel therapeutics.

## Introduction

The disruption of the methyl-CpG-binding protein 2 (MECP2), encoded on the X chromosome, is known to cause a severe neurodevelopmental disease called Rett Syndrome[1]. It is an X-linked dominant disorder observed mostly in female heterozygotes, as males with this mutation on their only X chromosome typically fail to survive birth[1]. Female patients with Rett Syndrome present with short stature overall, but have a relatively more profound microcephaly phenotype, suggesting a prominent role for MECP2 in the brain[2]. While the MECP2 protein is present in all tissues, the expression of MECP2 is particularly high in all types of mature neurons of the CNS[3]. Brain-specific deletion of MECP2 in mice phenocopies MECP2-null animals[4–7], and subsequent studies showed that neuronal specific deletion of MECP2 recapitulated phenotypes of MECP2-null animals.

MECP2 has been described as both a transcriptional stimulator and inhibitor, a regulator of RNA splicing, and a regulator of DNA methylation or reader of methylation[8–10]. Moreover, loss of MECP2 has been argued to lead to a variety of cell physiological defects such as mitochondrial permeability, diminished dendritic branching, altered electrophysiological activity, and changes in nuclear and nucleolar size, etc[11, 12]. Despite all these interesting observations, it is still not clear which of these are relevant for human Rett Syndrome patients, and which should serve as proxies for experimentally targeting and developing therapeutic strategies. The vast majority of these observations have been made *in vitro* with cell-based models of the disease or in murine models of the syndrome lacking MECP2[12], instead of in actual Rett patients. Therefore, the identification of which of these phenotypes occur in human patient brain is critical to understand the etiology of the disease and to uncover which *in vitro* or murine models most accurately reflect the human condition.

We previously created an isogenic *in vitro* system to model how the loss of MECP2 impacts development of human neural cell types by exploiting reprogramming of patient fibroblasts to a pluripotent state to create human induced pluripotent stem cells (hiPSCs), and differentiation towards particular neural lineages (Neural Progenitor Cells (NPCs) and interneurons)[13]. Our previously published work with this *in vitro* model of Rett Syndrome indicated that specifically postmitotic neurons lacking MECP2 show defects in dendritic branching coincident with induction of p53 and cellular senescence. Blocking senescence with P53 inhibition restored dendritic branching in Rett-patient-derived neurons[13]. These results were consistent with those of the Galderisi group who also showed senescence phenotypes in various Rett models[14, 15].

The study of the human Rett brain until very recently was limited to pathological examination of tissue or bulk molecular methods to understand the course of the disease. For example, Gogliotti et al performed bulk RNA-seq on a number of Rett brain samples versus unaffected controls, and found 100s of differentially expressed genes. However, the methods employed precluded identification of cell type specific changes and are complicated by the fact that the female Rett brain is chimeric for both MECP2 WT and MECP2 mutant cells based on random X-inactivation during development. A more recent study took advantage of differential SNPs between maternal and paternal alleles coupled with Single Cell-RNA-seq (sc-RNA-seq) to define gene expression changes in specific cell types[16]. While this study did identify differentially expressed genes in MECP2-cells, the analysis did not implicate particular dysfunctional pathways to shed light on the effect of loss of MECP2 in neurons.

Here we employ single-nuclei sequencing (sn-RNA-seq) on neurons from Rett brains and unaffected control brains to understand the effects of loss of MECP2. In addition, we re-analyzed data from previous Rett brain studies, as well as various *in vitro* models, to corroborate our findings and identify effective models of Rett Syndrome. We determined that while loss of *MECP2* leads to differential synaptic and metabolic gene expression patterns, which are reminiscent of neurons in aging brain, the ratio of subtypes of neurons within Rett brain is not dramatically altered. We also identify *POU2F1/OCT1* as a potential key transcription factor regulator of the stress response observed in *MECP2*-null neurons, and demonstrate that some of the key phenotypes found in Rett brain are recapitulated in *in vitro* models of the disease.

## Results

### Method to isolate and profile neurons with and without expression of MECP2 from individual Rett patient brains

Rett syndrome is caused by a heterozygous mutation in MECP2 gene. MECP2 is an X-linked gene, thus half of MECP2 alleles are turned off in a female brain due to random X chromosome inactivation, resulting in roughly half of the expressed alleles being wild type and the other half harboring MECP2 mutations[1, 17]. By isolating cells from a single brain, we can directly compare wild type cells to the mutant cells, without confounding variables such as genetic background or environment.

We used banked frozen brain sections of Rett patients to investigate differences in gene expression between MECP2^hi^ and MECP2^low^ cells. We took advantage of post-mortem tissue where we previously showed that Rett patient neurons harbor chimerism for MECP2 expression[13]. To preserve maximum genetic material following tissue thawing, we isolated the nuclei from these sections (Fig 1A)[18]. Nuclei isolation was performed utilizing sucrose gradient accompanied by a series of washes and centrifugations. The quality of nuclei isolation was assessed visually and quantitatively using a hemocytometer. We were particularly interested in the role of MECP2 in neurons because deletion of this protein in neurons has a particularly strong effect in Rett models, and because MECP2 is known to be expressed at a higher level in neurons than other cell types in the brain. Thus, we chose NeuN, a commonly used mature neuron nuclear marker to isolate neuronal populations from Rett nuclei. We stained the nuclei with MECP2, NeuN and DAPI and subjected them to FACS. We collected DAPI^+^/Neun^+^ intact neuronal nuclei, which were either MECP2^hi^ or MECP2^low^. To confirm the nuclei sorting strategy was successful, we used cytospin, immunostaining, and western blotting which confirmed that the expression level of MECP2 was highly diminished in the MECP2l^ow^ population of neurons (Fig 1B and C).

**Figure 1.**
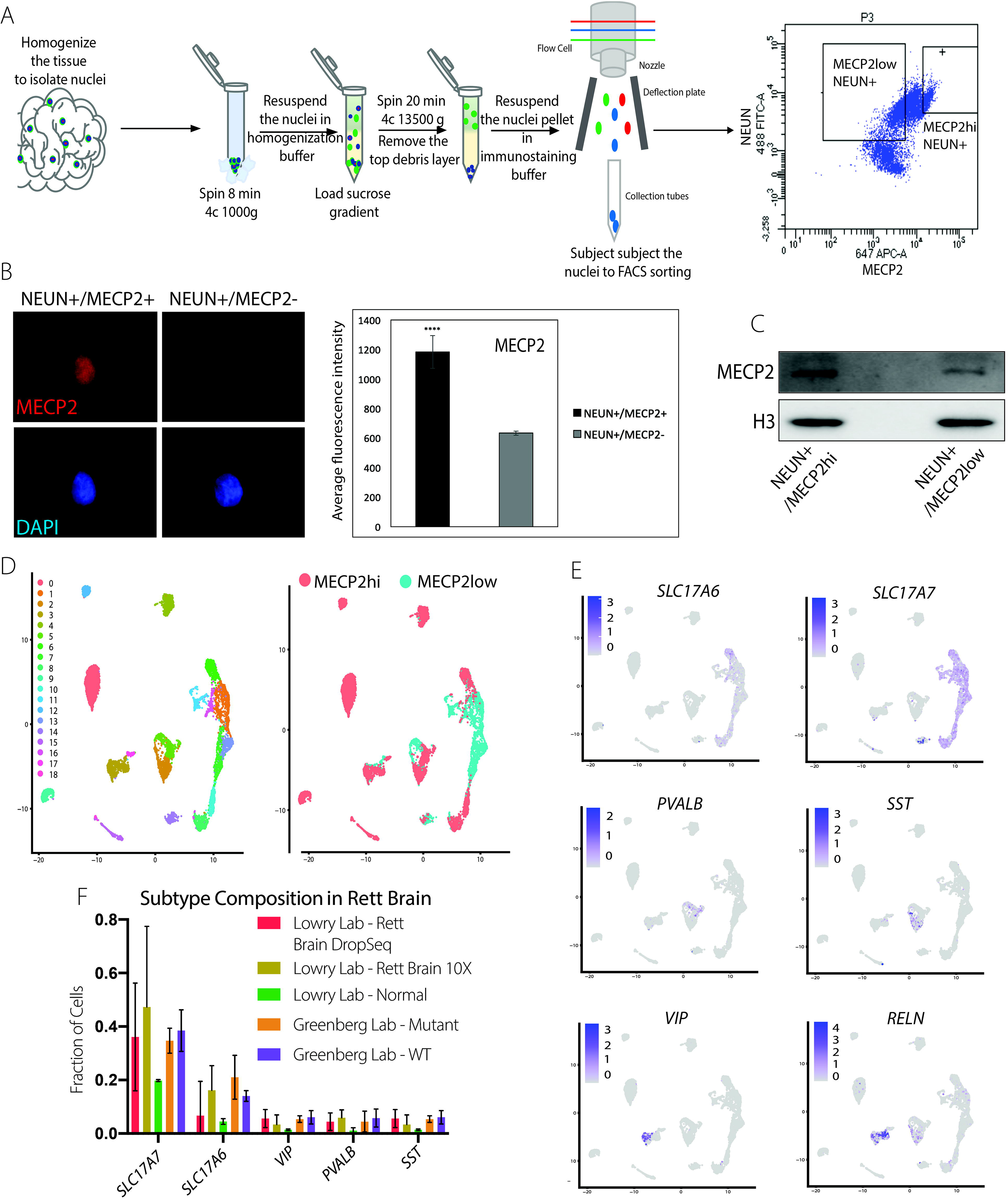
A method to isolate and molecularly profile MECP2^hi^ and MECP2^low^ neurons from Rett brain. (**A**) The diagram depicts the process of nuclei isolation that employs sucrose gradient and a number of washes and centrifugations followed by immunostaining and FACS. (**B**) Immunostaining of the sorted MECP2^hi^ and MECP2^low^ neuronal nuclei after cytospin from Rett patient brain shows a decrease in MECP2 expression in the MECP2^low^ fraction, images taken at 40x. (right), quantification of IF signal. Statistical significance was checked using a paired t-test, where * p < 0.05, ** p < 0.01, *** p < 0.001, **** p < 0.0001. (**C**) Western blot analysis verifying a decrease in MECP2 expression in the MECP2^low^ fraction isolated by FACS. (**D**) UMAP showing 19 clusters identified during the analysis of RNA-seq. MECP2^low^ (MUT) and MECP2^hi^ (WT) nuclei contribute to multiple identified clusters, expressing excitatory and inhibitory neuronal subtypes. (**E**) UMAP plots of abundant neuronal subtypes as defined by the indicated markers. (**F**) Proportions of neuronal subtypes in MECP2^hi^, MECP2^low^ and WT nuclei from three independent experiments. Mutant (MECP2-Mutant nuclei from Rett brain), WT (MECP2-wild type nuclei from Rett brain), Normal (wild type nuclei from wild type brain).

Wild type and mutant neuronal nuclei were separately analyzed by both Drop-SEQ (3 individual Rett patients)[19] and 10x RNA sequencing (10x Genomics) (2 individual Rett patients). While both methods detected numerous clusters related to various neuronal subtypes and were largely concordant with one another, the coverage of the 10X capture was more substantial and thus became the focus of our follow-on analyses (Supplemental Figure 1). The drop-seq data were sufficient to identify neuronal subtypes and were thus used to explore cell proportion changes in response to loss of MECP2. Clustering of the 10X data generated 19 transcriptionally unique clusters in both the WT and mutant populations. Further analysis of the gene expression revealed several subtypes each of inhibitory and excitatory neuronal populations (Fig 1D).

This sorting and transcriptomic approach allowed for a determination of whether the proportion of cell types was affected by MECP2 expression. Because NeuN expression tracked nearly identically with MECP2 expression in wildtype neurons (not shown), and because all identified subtypes of neurons expressed NeuN and MECP2 to a similar degree, we were able to construct a proportional analysis of cells types between MECP2^hi^ and MECP2^low^ neurons. In total, we generated data from 5 individual brains (with both Drop-seq and 10X), and we also re-analyzed data from Renthal et al that used SNP analysis to define WT vs mutant neurons from the Rett brain. Taken together, this analysis from 8 Rett samples did not reveal a significant difference in the ratio of any neuronal subtypes amongst all these Rett patient brains (Fig 1E). These data suggest that neurological deficits in Rett patients are probably not due to different proportions of neurons.

We next performed differential expression analysis and applied a *p-value* cut off of 0.01 to identify differentially expressed genes (DEGs) between MECP2^hi^ and MECP2^low^ neurons from each brain. We first compared DEGs identified in our own data versus those uncovered from Renthal et al to identify overlap between the two methods (Supplemental Figure 2). In fact, DEGs from our 10X analysis overlapped significantly with data from the Greenberg group with independent brain samples despite the differences in methods (identification of WT vs Mutant and sc-RNA-seq approaches) (Figure 2A), indicative of the robustness of our approach. We next showed that there are overlapping DEGs patterns found in both excitatory and inhibitory neurons in Rett brain (Figure 2B), suggesting general physiological defects due to loss of MECP2. Looking at DEGs across various neuronal subtypes from our own data, we noticed that only a subset of neurons appear to be affected transcriptionally by loss of MECP2, suggesting there is something of a selective vulnerability (Figure 2C) as only particular subtypes of neurons showed significant transcriptional effects to loss of MECP2 despite the fact that this protein is ubiquitously expressed by all neurons. Another surprising finding was the BDNF, a gene that has been proposed to be a “target” of MECP2 regulation in scores of studies was only found to be strongly expressed in one excitatory neuron cluster (SLC17A7+/NR4A2+), and was not differentially expressed between MECP2^hi^ and MECP2^low^ neurons. While this is consistent with the Renthal et al study[16], this is in contrast to numerous studies in murine and human models of Rett Syndrome over the last 15 years, and could highlight the importance of studies of authentic disease patient tissue.

**Figure 2.**
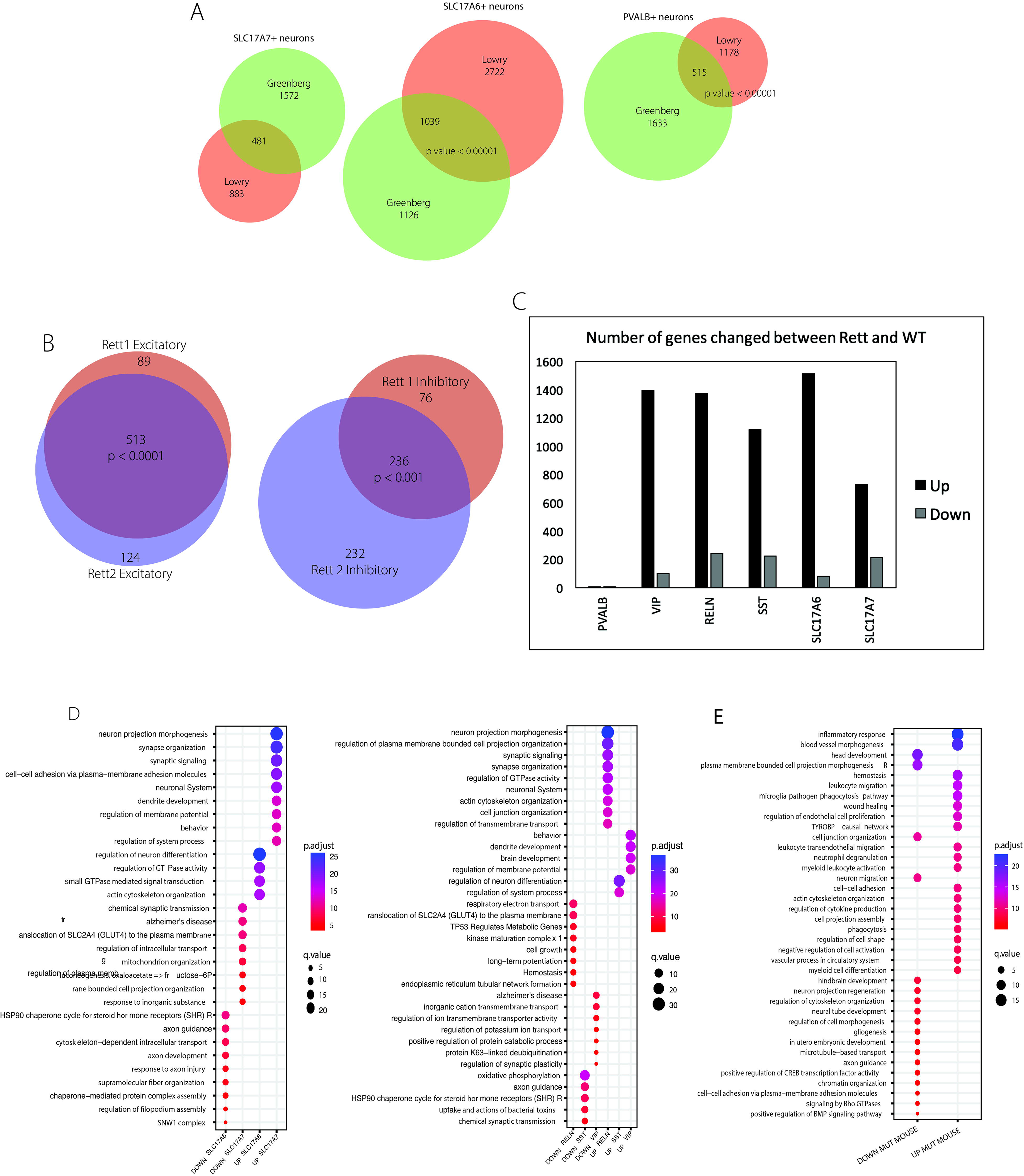
Transcriptome analysis reveals alteration in synaptic and metabolic gene expression due to lack of MECP2. (**A**) Venn diagram showing the number of overlapping genes in several excitatory and inhibitory neuronal populations between our data and a published data set from the Greenberg lab. (**B**) Venn diagram showing the number of genes that overlap between our excitatory and inhibitory neuronal populations. (**C**) A summary of the number of Differentially Expressed Genes (DEGs) in several of the main cell types. (**D**) Gene ontology (GO) analysis of misregulated pathways indicate significant alterations in synaptic and metabolic gene expression in both excitatory (left) and inhibitory (right) MECP2^low^ neurons. (**E**) GO analysis of sc-RNA-seq data from MECP2 null transgenic brain neurons.

We then used Gene Ontological (GO) analysis to determine which patterns of gene expression are upregulated and downregulated in either excitatory or inhibitory subclasses of neurons in the absence of MECP2. Remarkably, in the two major classes of excitatory neurons (SLC17A6+ and SCL17A7+), nearly all the gene expression changes induced in MECP2^low^ neurons appeared to be related specifically to neuronal physiology such as ‘Neuron Projection Morphogenesis’, ‘Synapse Organization’, ‘Dendrite Development’, ‘Neuron Differentiation’ etc. This is an important observation considering recent data from our own group showing that Rett patient derived neural organoids show elevated levels of synapses and synaptic genes (Samarsinghe et al, 2021). On the other hand, the majority of gene expression changes downregulated, appeared to be related to cellular metabolism, such as ‘Translocation of GLUT4 to membrane’, ‘Mitochondrion Organization’, ‘Gluconeogenesis’ etc. Considering the overlapping pattern of DEGs shown in Figure 2B, it is not surprising that the same GO analysis on inhibitory neuron subtypes (RELN+, SST+, VIP+) yielded similar results, where the same categories of DEGs were found in both the up- and downregulated expression patterns. While it is perhaps not surprising that genes related to neuronal physiology were perturbed in MECP2^low^ neurons in the Rett brain, considering previously reported phenotypes from both *in vivo* and *in vitro* phenotypes, the alteration of metabolism in Rett brain is relatively unexplored and warranted further investigation. This pattern of metabolic disruption at the RNA level could portend metabolic defects in the neurons, as it has been previously established that metabolic dysfunction can present at the level of RNA expression of important metabolic enzymes[20, 21].

Numerous attempts to model Rett Syndrome have taken advantage of deletion of Mecp2 via homologous recombination[4, 5, 22–30]. These models recapitulate some, but not all phenotypes of human Rett Syndrome. We re-analyzed a dataset from murine Rett brain and found first that many of the same neuronal-specific categories of genes were downregulated instead of upregulated as in the human Rett brain, and that most of the upregulated genes were related to inflammatory responses such as ‘Inflammatory Response’, ‘Leukocyte Migration’, ‘Wound Healing’ etc (Figure 2E). It is tempting to speculate that the differences observed between the human and murine cases is due to the design of the mouse model, whereby Mecp2 was deleted as opposed to mutated, which could have induced a more profound phenotype than that observed in human.

Next, we looked at the published RNA sequencing data sets that recently became available for other syndromes such as Alzheimers, Aging, and Autism-Spectrum Disorders. Here we demonstrate that only the Aging brain showed a similar pattern to the Rett brain whereby genes related to synapse and metabolism pathways were misregulated. The Aging brain also showed a strong induction of inflammatory gene signatures similar to what we observed in the murine Rett brain such as ‘Inflammatory Response’, ‘Positive Regulation of NFKB’, ‘Microglial Activation’ etc. Conversely, downregulated gene categories were almost entirely related to metabolism such as ‘Mitochondrion Organization’, ‘Inner Mitochondrion Membrane Organization’ and ‘Aerobic Respiration’, nearly identical to what was observed in the human Rett brain. (Supplemental Figure 3) These data suggest that recent evidence for a role in senescence and neuronal stress in Rett neurons could be akin to premature aging of the neurons in Rett patients[13–15, 31–36].

### POU2F1/OCT1 binding sites are enriched in genes affected by loss of MECP2 function

To gain more insight into the molecular nature of the changes in the Rett patient brain, we analyzed transcription factor binding sites enriched within the promoter of significantly altered genes (Figure 3A). A similar pattern emerged from analysis of the Greenberg Rett sc-RNA-seq data (Figure 3C), as well as in the Gogliotti bulk RNA-seq data from Rett brain (Figure 3D). Furthermore, looking across the Aging brain, Rett Brain, and *in vitro* culture, in every case, POU2F1/OCT1 binding sites were also among the most enriched in both up and downregulated genes (Figure 3, and 4). This suggests that POU2F1/OCT1 could be playing an outsized role in neuronal aging and in the response of neurons to loss of MECP2 in a variety of settings. This finding is interesting in light of extensive data suggesting that POU2F1/OCT1 is important in mediating stress responses in a variety of tissues[37–40], and the previous work from our group and the Galderisi group that MECP2^low^ neurons undergo upregulation of stress response pathways downstream of P53[13–15].

**Figure 3.**
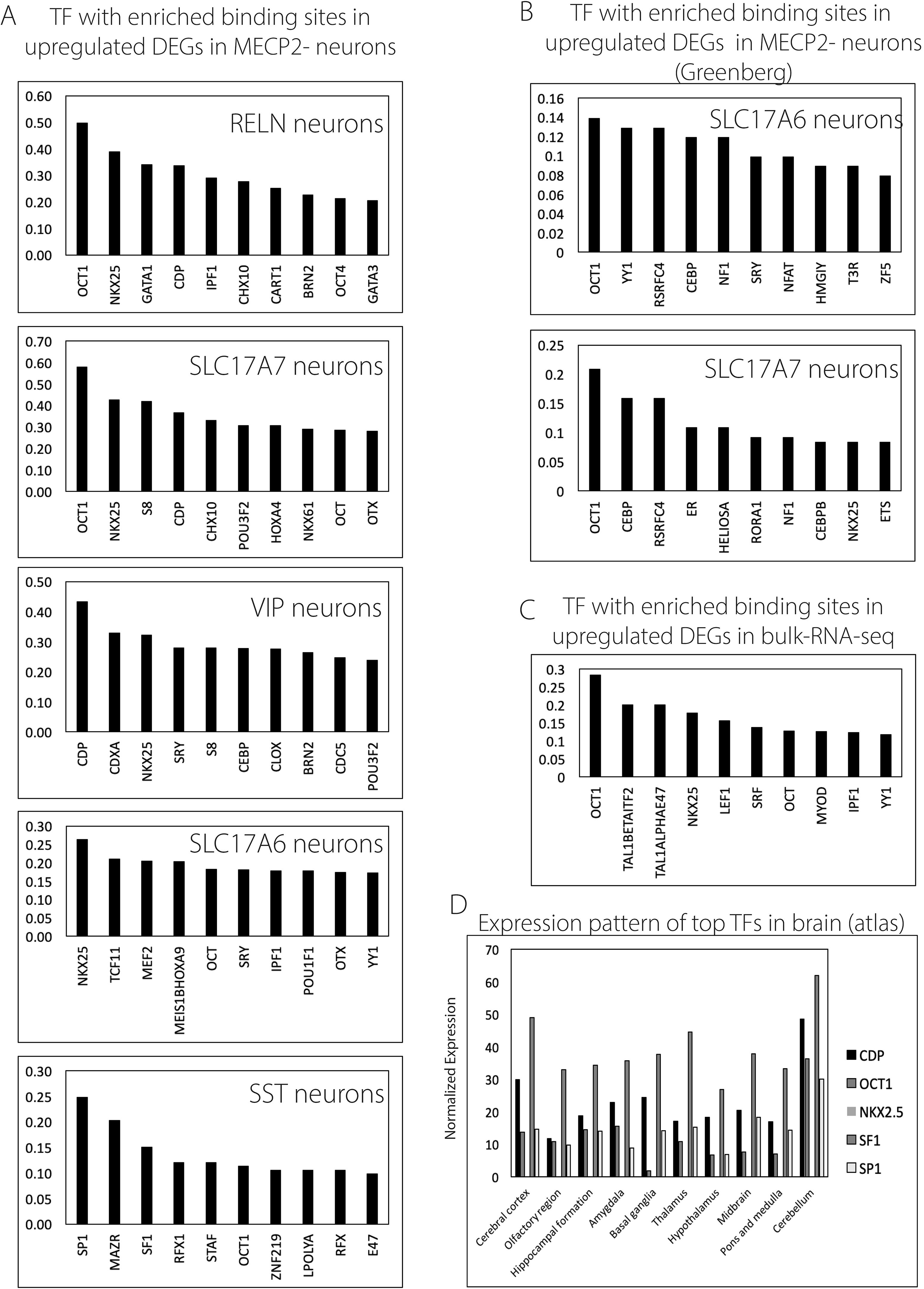
POU2F1/OCT1 is one of the most enriched potential regulators of genes altered in Rett patient brain MECP2^low^ neurons. (**A**) Analysis of the transcription factor binding sites within the genes significantly altered dues to lack of MECP2 shows POU2F1/OCT1 binding site enrichment (Measured by DIRE algorithm). y-axis indicates the percentage of genes containing a conserved binding site for a particular transcription factor. (**B**) Same analysis as in (**A**), but using the Greenberg Rett dataset. (**C**) Same analysis as in (A), but performed on bulk-RNA-seq DEGs from Gogliotti et al. (**D**) Expression of the enriched potential regulators of genes altered in MECP2^low^ neurons by brain region. Analysis of Rett patient derived *in vitro* neurons shows enrichment of POU2F1/OCT1 binding sites within the differentially expressed genes.

**Figure 4.**
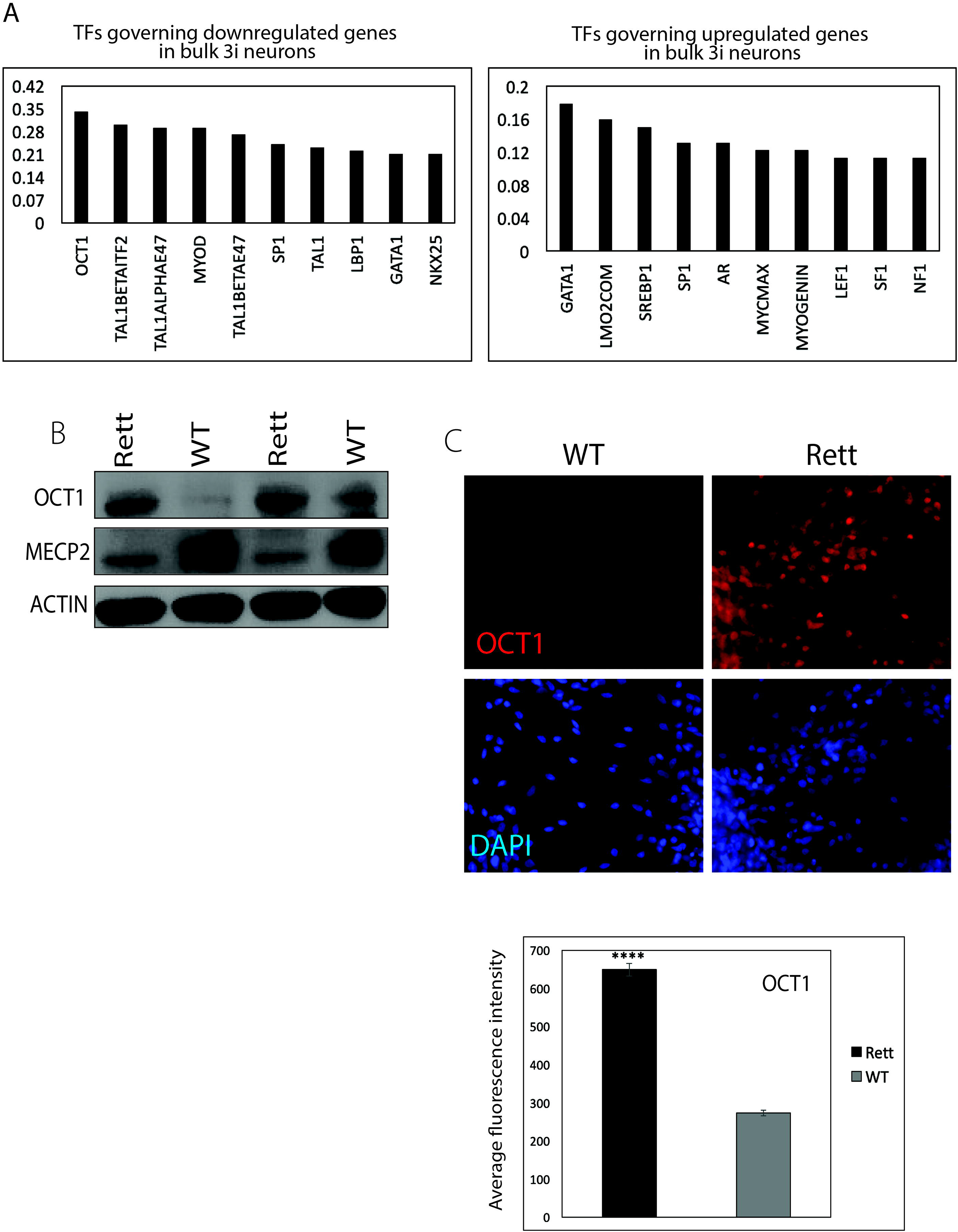
POU2F1/OCT1 expression is induced in neurons lacking MECP2 *in vitro*. (**A**) DIRE analysis on RNA-seq data from Rett neurons in vitro showed enrichment for OCT1 binding sites in DEGs. (**B**) Left, immunostaining shows significantly higher expression of POU2F1/OCT1 in patient derived Rett neurons compared to the wild type, 40x. Right, quantification of the immunostaining. (**C**) Western blot shows that POU2F1/OCT1 is induced in two independently derived Rett cell lines compared to two wild types.

To corroborate our findings and identify faithful *in vitro* models of Rett Syndrome, we investigated whether similar gene expression patterns are present in a cell culture model of Rett. We analyzed bulk RNA sequencing results of interneurons generated in our lab through a standard protocols using a combination of growth factors and three pharmacological inhibitors of signaling pathways[41, 42]. As in the Rett brain neurons, of *MECP2 in vitro* caused downregulation of ‘Axon development’, ‘Synaptic signaling’, ‘Synapse assembly’ and ‘Cerebellum development’. In addition, MECP2^low^ neurons showed induction of stress pathways such as ‘Wound healing’ and ‘Negative regulation of cell proliferation’, similar to what was seen in Aging brains. Comparing isogenic neurons both wild type to mutant for *MECP2* expression revealed that *POU2F1/OCT1* was significantly upregulated at the protein level in the absence of *MECP2* by western blotting and immunofluorescence (Figure 4A, B).

### Potential routes to activation of POU2F1/OCT1

POU2F1/OCT1 is known to be regulated post-translationally through covalent modification and stabilization from degradation [43, 44]. Several studies have suggested that DNA damage or lack of DNA repair is a stressor sufficient to stabilize POU2F1/OCT1 levels in the cell. Our data from Rett brain suggests that metabolic disruption is present and this could be another route to stabilization of POU2F1/OCT1. Therefore, to determine what types of stress in human neurons can stabilize POU2F1/OCT1 protein, we treated wildtype human neurons with various stress inducing insults and measured POU2F1/OCT1 protein levels. Because the sc-RNA-seq data from Rett brains showed evidence of neuronal dysfunction due to metabolic stress, we attempted to see which of these could potentially be relevant in *in vitro* models of Rett Syndrome, and which could potentially induce POU2F1/OCT1 as a result.

In Ohashi et al[13] we showed evidence that MECP2^low^ neurons potentially have increased DNA damage by staining for H2AX, even in the absence of deliberate DNA damage[13] using an interneuron differentiation scheme. Here, we repeated this type of study using neurons generated by growth factor withdrawal to determine the general applicability of these findings. We found that activation ofATR was significantly increased upon UV induced DNA damage, and that MECP2^+^ and MECP2^−^ neurons responded similarly to the damage (Figure 5A). In addition, we found that inducing DNA damage in WT neurons led to a strong increase in POU2F1/OCT1 protein expression (Figure 5A and B), possibly indicating a role for POU2F1/OCT1 as a stress sensor also in human neurons.

**Figure 5.**
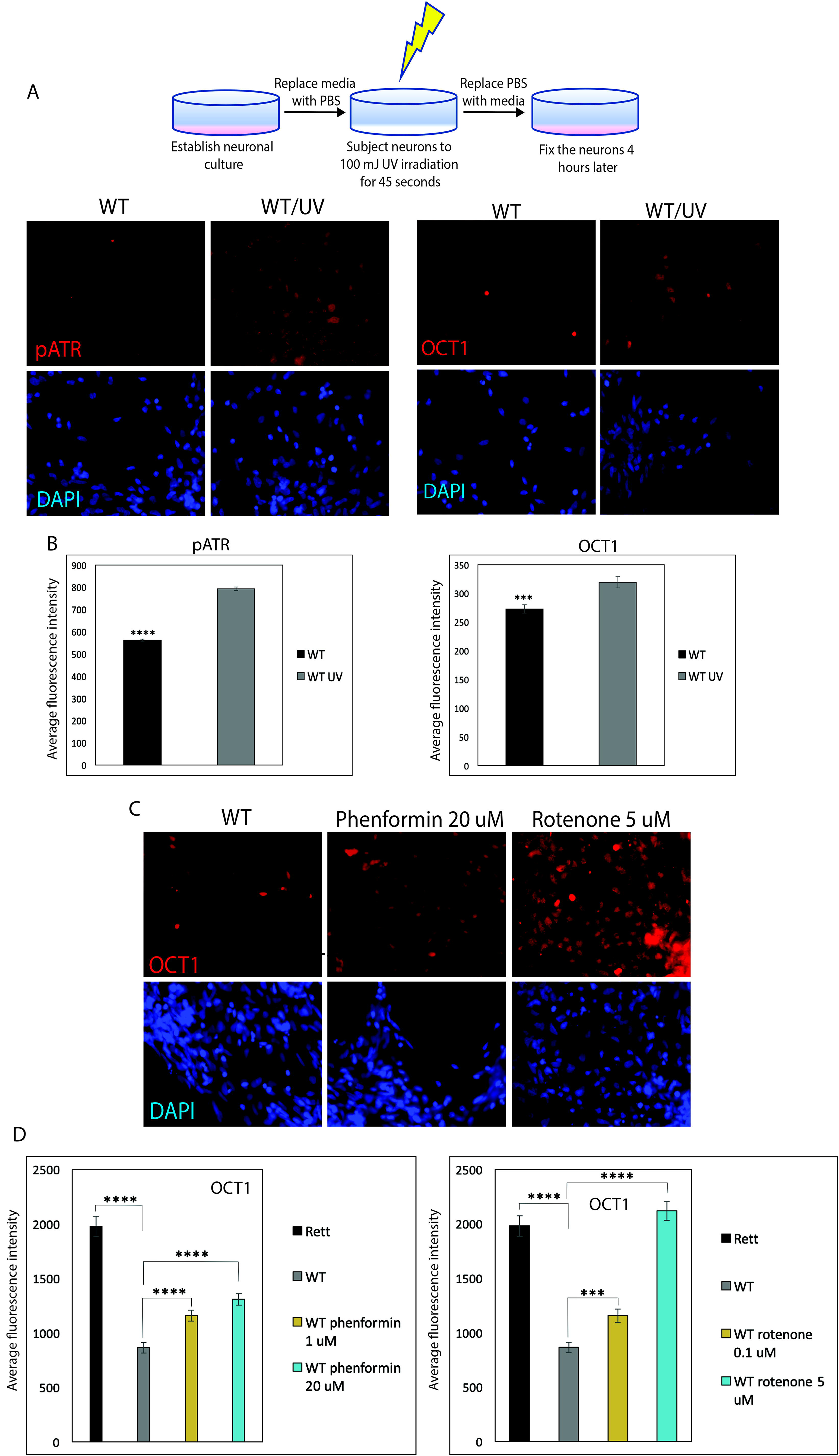
POU2F1/OCT1 is induced in wild type neurons upon DNA damage and disruption of mitochondrial function *in vitro*. (**A**) Left, schematic of UV irradiation of wild type neurons. Middle, immunostaining showing that phosphorylation of ATR increases in neurons upon UV irradiation, indicating DNA damage, 40x. Right, quantification of the immunostaining. (**B**) Left, immunostaining showing that Induction of DNA damage in wild type neurons leads to elevated expression of POU2F1/OCT1 protein, 40x. Right, quantification of the immunostaining (**C**) Immunostaining showing that wild type neurons treated with 20 uM Phenformin or 5 uM Rotenone which leads to a disruption of mitochondrial function, induce of POU2F1/OCT1 expression, 40x. (**D**) Quantification of the immunostaining, POU2F1/OCT1 is induced upon Phenformin or Rotenone treatment in a concentration dependent manner.

Mitochondrial dysfunction is known to lead to induction of stress pathways such as P53[45]. Therefore, we attempted to disrupt mitochondrial function *in vitro* to ascertain whether metabolic stress could induce POU2F1/OCT1 protein stabilization in human neurons, as we predict is happening in Rett neurons *in vivo*. In fact, disruption of metabolic homeostasis through the use of uncoupling agents or inhibitors of the electron transport chain led to a significant induction of POU2F1/OCT1 protein in otherwise normal neurons to nearly the level observed in Rett mutant neurons (Figure 5C, D). Taken together, these experiments in Figure 5 suggest that either metabolic aberration or activation of DNA repair pathways in the Rett brain could potentially be driving changes in POU2F1/OCT1 activity or expression that then affect the expression of numerous target genes as reflected in the sc-RNA-seq data.

Based on the ontological analysis of the transcriptional data from the Rett brain, we hypothesized that glucose metabolism and pyruvate oxidation pathways could be disrupted in Rett neurons. To test this hypothesis *in vitro*, we next used glucose tracing and metabolomics by Mass Spectrometry to assess the utilization of glucose in neurons with and without MECP2 function, and found defects in several TCA metabolites, which would be consistent with mitochondrial dysfunction due to lack of pyruvate oxidation and energy production through the electron transport chain (Figure 6A). Repeated metabolomics experiments were compiled and organized into pathway enrichment scores for the observed metabolic changes (similar to Gene Set Enrichment Analysis (GSEA) for RNA data) leading to categories such as ‘Pyruvate metabolism’ (Figure 6B). To functionally measure whether Rett neurons lacking MECP2 expression show defective cellular metabolism, we used Seahorse assay to determine their oxygen consumption rate, a direct output of mitochondrial function, particularly the TCA cycle and the electron transport chain, which was predicted to be affected in Rett neurons both in patients transcriptionally and in our i*n vitro* model via metabolomics. The Seahorse assay showed that Rett mutant neurons from two distinct Rett patients are defective at all stages of the electron transport chain (Figure 6C). Further analysis of Seahorse data from these experiments show that these same Rett neurons from two patients had a reduced ATP consumption rate relative to isogenic neurons expressing MECP2 (Figure 6D).

**Figure 6.**
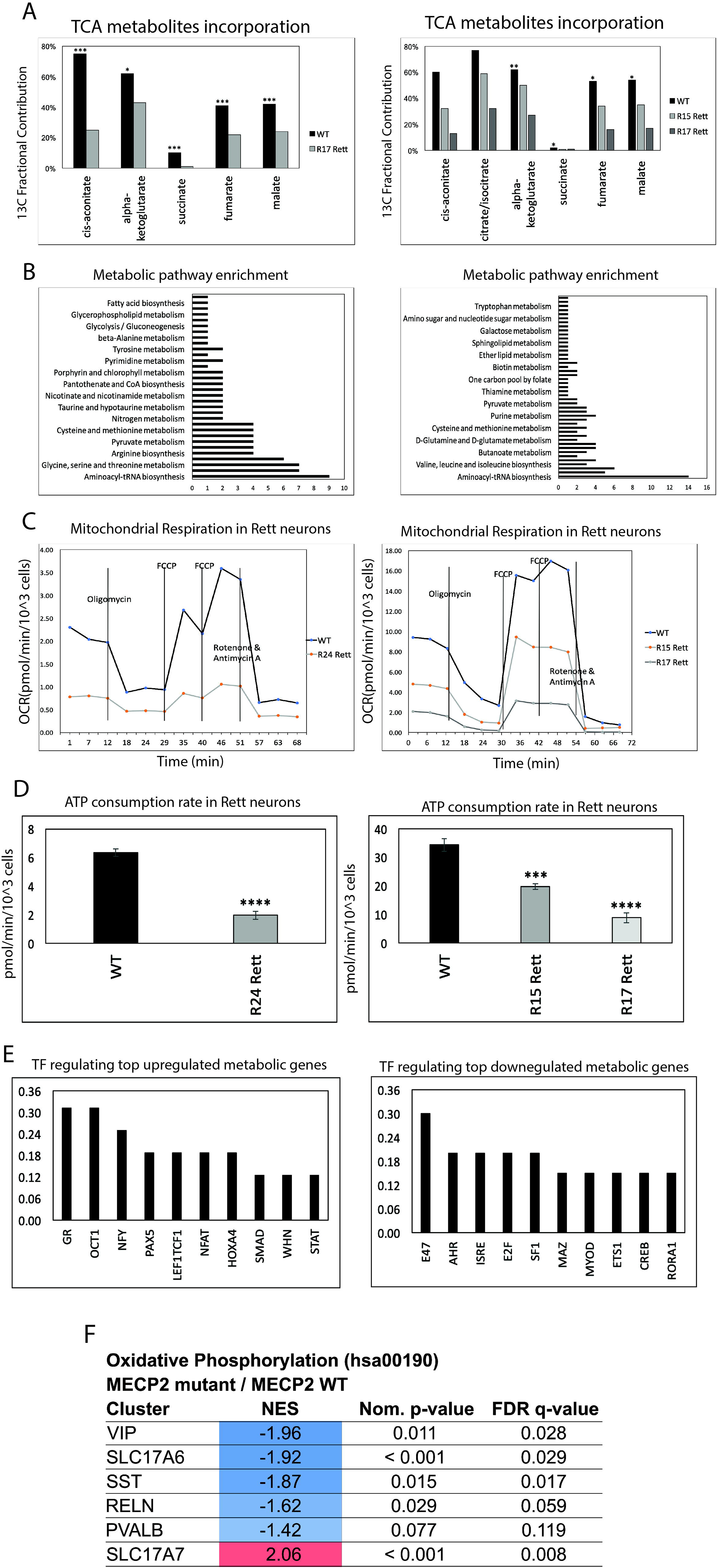
Rett neurons exhibit metabolic dysfunction *in vitro*. (**A**) Lack of MECP2 in patient derived neurons leads to a decreased incorporation of C13 labeled glucose into the citric acid cycle. (**B**) Pathway enrichment showing the processes most affected by the metabolic dysfunction in Rett neurons. x-axis is the number of times a particular pathway came up as affected during the analysis. (**C**) Oxygen consumption rate is decreased in Rett neurons as detected by the Seahorse assay. (**D**) Rett neurons also exhibit decreased ATP consumption rate. (**E**) Analysis of transcription factor binding sites within significantly changed metabolic genes revealed POU2F1/OCT1 as being one of the potential regulators of these genes. (**F**) GSEA analysis of Rett neurons from patient brain predicts diminished activity of TCA cycle in most identifiable neuronal subtypes in Rett brain.

We also reanalyzed the DEGs shown in Figure 3 to isolate just those called out as ‘Metabolism related’ by gene ontology, and used transcription factor binding site enrichment analysis to identify transcription factors that potentially regulate differences in metabolic gene expression. In the Metabolic genes upregulated, POU2F1/OCT1 again stood out as a potential regulator of these genes (Figure 6E). To take an unbiased approach to analyze metabolic pathways in the Rett brain, we next used Gene Set Enrichment Analysis (GSEA)[46] across all metabolic pathways from KEGG[47]. This method allowed for comparison of the relative metabolic pathway expression in different neuronal cell types in the brain. This analysis demonstrated that most neuronal subtypes in the Rett brain have significantly reduced expression of the oxidative phosphorylation pathway (Figure 6F), consistent with the reduced TCA cycle activity observed in our *in vitro* model (Figure 6A-C). Together, these data are consistent with mitochondrial dysfunction in the absence of MECP2 function, which perhaps sheds light on some of the transcriptional differences observed in Rett brain. As others have already identified metabolic deficiencies in murine and human Rett experimental models[31, 32, 48], the single cell analysis performed here on human Rett brain suggests a similar dysfunction potentially driving Rett phenotypes in patients.

## Discussion

This study sheds new light on the etiology of Rett Syndrome by molecular interrogation of individual neurons from Rett patient brain tissue. From these data, several important observations were made about selective susceptibility of individual subtypes of neurons, the proportions of neuronal subtypes, effects on expression of synaptic genes, and the nature of stress in neurons induced by loss of *MECP2* function. We identified a transcription factor that potentially plays a role in regulating the differential gene expression observed in some subtypes of neurons lacking *MECP2* function. We used *in vitro* models to both confirm what we had observed in Rett brains, but also to extend this work and determine the nature of the stress

Given their critical role in cognition, it is especially important that neurons be equipped to withstand various threats such as injury and disease. This is especially true for postmitotic cells, which bear special relevance to Rett Syndrome because MECP2 is most highly expressed in these specifically [6, 7, 49]. It is known that many forms of neuronal stress response lead ultimately to the activation of various transcription factors, which in turn act to determine cell fate[50–52]. Some hypothesize that the resulting transcriptional changes may be a cellular attempt to mitigate the initial stress or its downstream impacts, remove damaged neurons, alter brain activity to compensate for the stressor, or may even directly be a pathological symptom [53]. Further, neuroinflammation, a known consequence of chronic stress [54], has been linked to MECP2 mutations as measured by observed dysregulation of several acute-phase response (APR) proteins in a Rett mouse model [55]. The current study attempted to define which types of neurons in the Rett brain are undergoing stress and what are potential molecular or physiological triggers of that stress.

Studies from our group and others have suggested an important role of senescence in Rett Syndrome. It has been argued that senescence may be a cellular adaptation designed to conserve energy required for division and differentiation so the cell can survive when facing stress[56]. The Galderisi group has published several studies demonstrating senescence in Rett patient mesenchymal stem cells (MSCs) [14, 15], partially MECP2 silenced human MSCs[14], MECP2 silenced human neuroblastoma cells [32], and heterozygous MECP2 mutant mouse mesenchymal stromal cells [31] and neural stem cells [32]. In our previous study on Rett derived hiPSCs, we found SASP induction in MECP2 null interneurons, consistent with these previous results[13]. While neurons from the human Rett brain did not show clear transcriptional evidence of a senescence program, there was evidence for metabolic stress.

Given their critical role in cellular energy production, mitochondrial function is highly important for normal cell function. We found strong evidence for mitochondrial disruption in the Rett brain via ontological analysis of differentially expressed genes (Fig 6). These data were consistent with an increasingly large body of evidence which connects Mecp2 deficiency to mitochondrial abnormalities, and in turn these abnormalities to further Rett phenotypes. Multiple studies in Rett mice models and human Rett samples have shown that Mecp2 interacts with genes encoding mitochondrial subunits which are abnormally expressed in its absence [57–59]. Additionally, analysis of male mice hemizygous for Mecp2 mutation revealed redox-imbalance across cytosol and mitochondria of neurons accompanied by increased oxygen consumption and generation of reactive oxygen species (ROS) [34]. Increased oxygen consumption was also confirmed in NPCs derived from Rett Syndrome patient cells in a separate study[60]. This is particularly interesting since another characteristic Rett mitochondrial alteration is a leaky inner membrane [61].

Our data suggest that MECP2^low^ neurons have dysfunctional mitochondrial metabolism, which is consistent with previous studies. This is interesting in light of what is known about mitochondrial diseases caused by mutations in genes for mitochondrial proteins. These diseases share many of the characteristic clinical symptoms of Rett Syndrome including intellectual disability, motor problems, and seizures [62, 63]. In addition, they are accompanied by evidence of oxidative damage and increased blood lactate and pyruvate content [64] which have also been identified as features of Rett [34, 61, 65].

There is also extensive evidence of the role of abnormal mitochondria in oxidative stress, a type of neuronal stress heavily implicated in many diseases including Rett Syndrome which leads to the before mentioned oxidative damage. Oxidative stress is a cellular condition caused by excessive ROS. This therefore suggests a positive feedback mechanism in which mitochondrial defects (initially stemming from MECP2 loss) lead to overproduction of ROS which in turn causes further mitochondrial defects due to oxidative stress. This cycle may explain the delayed onset of Rett as the initial mitochondrial deficiency caused by loss of MECP2 may not produce a large enough effect to lead to symptoms, but accumulation of mitochondrial defects over time can. However, this hypothesis conflicts with the fact that unlike many other neurological diseases in which oxidative stress is implicated (such as Alzheimer’s and Parkinson’s) [53], Rett Syndrome is not degenerative. Additionally, reintroduction of Mecp2 into both Mecp2 null and deficient symptomatic mice models rescued wildtype levels of several oxidative stress markers[5, 66]. One possible explanation for this is that once the initial driver of mitochondrial dysfunction is ameliorated, the brain is able to correct for the accumulation of mitochondrial defects, likely through the major mtDNA repair mechanism, base excision repair. However, further studies are required to determine if this is true, and more generally how the dynamics of cellular stress and cellular stress response work together to produce Rett symptoms without typical neurodegeneration.

Critically, oxidative stress is known to induce senescence, p53 activation, and Oct1 upregulation, both of which we have identified as key features of MECP2^low^ neurons. The Oct family, one branch of the POU transcription factors, is a group of transcription factors so named for their characteristic recognition of the octamer DNA element ATGCAAAT. Unlike any of its family members, Oct1 is expressed ubiquitously both temporally and spatially throughout the body [37, 38]. A study on Oct1 deficient mice revealed abnormal expression of many stress related genes as well as hypersensitivity to oxidative stress[38, 39, 67]. Based on the observation that many of the genes dysregulated in response to stress inputs contained human-conserved octamer sequences in their regulatory regions, it is hypothesized that Oct1 may play an important role in regulation of cell-stress response genes. However, other findings suggest that Oct1 may be a direct participant in cellular response pathways[37]. Oct1 has been shown through various studies to interact with multiple stress-response implicated factors. One of these studies found that Oct1 mediates the induction by BRCA1 of the gene GADD45[68]. This is of particular interest since GADD45 is a stress-inducible DNA repair gene regulated by p53[69]. In addition, we found that GADD45 was one of several p53 targets whose expression was upregulated following silencing of MECP2 in neuronal cultures.

Another question not answered by the current study is exactly how loss of MECP2 function can lead to metabolic dysfunction. While MECP2 is known to bind methylated DNA, it is also now clear that MECP2 can bind RNA as well and this protein has been implicated in a panoply of different functions. One could speculate that loss of function of an epigenetic regulator such as MECP2 could disrupt gene regulation of important metabolic proteins; or perhaps induce a stress response that leads to P53 upregulation and subsequent decrease of metabolic activity in an active effort to diminish ROS levels and allow for repair; or many have argued that MECP2 can regulate gene expression as a DNA binding protein or more recently through its ability to directly bind RNA. Finally, it is tempting to speculate that because epigenetic regulation and metabolism both share similar substrates and products, that perhaps epigenetic regulation in the nucleus leads to an imbalance of substrates available for metabolic enzymes, and therefore metabolic disruption.

Here we found that POU2F1/OCT1 is potentially a key regulator of gene expression in response to loss of MECP2 in Rett brain, and is clearly upregulated in MECP2-null neurons *in vitro*. Furthermore, we probed for potential instigators of OCT1 stabilization and found that metabolic stress and DNA damage can induce OCT1 in normal neurons. Whether OCT1 induction is required for the physiological manifestations associated with Rett disease in patients is an open question. While it does seem likely that OCT1 induction is due to increased stress in neurons lacking MECP2, it is not clear whether blocking OCT1 function in neurons would reverse synaptic transmission defects observed in other studies in response to loss of MECP2. It is also possible that increasing OCT1 further would promote the type of repair or abrogation of stress necessary to promote neurophysiological restoration of MECP2-null neurons. Future effort will be devoted to understanding how OCT1 functionally regulates neuronal stress in Rett brain.

## Materials and Methods

### Generation of *in vitro* neurons

First, isogenic Rett Syndrome and wild type iPSCs were derived from Rett patient fibroblasts as described previously [13]. iPSCs were maintained on plates coated with matrigel (Corning) in mTeSR1 (StemCell Technologies) until 80% confluency. Neural progenitor cell (NPC) fate was induced following the manufacturer’s protocol. Briefly, 2 million cells were passaged per each well using Accutase (StemCell Technologies) into the STEMdiff SMADi neural induction medium (StemCell Technologies). Cells were passaged this way once a week, two more times to induce NPC fate. Next, NPCs were differentiated to a Cajal-Retzius neuronal cell type using growth factor withdrawal method. EGF and FGF were removed from the media, and the cells were cultured in DMEMF12 supplemented with N2 and B27 (Thermo Fisher).

### Western Blot

Cell lysate was prepared using RIPA buffer (Pierce) supplemented with Halt Protease Inhibitor Cocktail (ThermoFisher Scientific) and Halt Phosphatase Inhibitor Cocktail (ThermoFisher Scientific). Total protein concentration was determined using BCA Protein Assay Kit (ThermoFisher Scientific) following the manufacturer’s protocol. Equal protein concentrations were loaded onto the NuPAGE 4-12% Bis-Tris gel (ThermoFisher Scientific) and run at 150 Volts for 90 minutes in running buffer, containing 25 mL of 20x NuPAGE MOPS SDS Running Buffer (ThermoFisher Scientific) and 475 mL of mili-Q water. Next, the protein was transferred onto the nitrocellulose membrane at 30 Volts for 60 minutes in transfer buffer, containing 25 mL 20x NuPAGE Transfer Buffer (ThermoFisher Scientific), 100 mL Methanol (ThermoFisher Scientific), 375 mL mili-Q water. The membrane was blocked overnight at 4°C in OneBlock Western-FL Blocking Buffer (Genesee Scientific), then incubated in the primary antibody at 4°C overnight. The following primary antibodies were used: rabbit MECP2 (Diagenode C15410052, 1:1000), rabbit anti-histone H3 (Abcam ab1791, 1:1000), rabbit OCT1 (Novus Biologicals NBP2-21584, 1:500), mouse ®-actin (SCBT sc-47778, 1:500). Membrane was washed twice with 0.1% PBST and incubated in anti-rabbit or anti-mouse secondary HRP-labeled secondary antibody (ThermoFisher Scientific 31460, 31430 1:100000) for 1 hour at room temperature. Membrane was washed twice with 0.1% PBST and SuperSignal West Femto Maximum Sensitivity Substrate (ThermoFisher Scientific) was added to the membrane and subjected to film exposure.

### Immunofluorescence and image quantification

Cells grown on coverslips were washed with PBS and fixed with 4% paraformaldehyde (Electron Microscopy Sciences) for 15 minutes at room temperature. Next, the cells were washed with 0.1 % PBST three times and blocked in MAXblock Blocking Medium (Active Motif) for 1 hour at room temperature, then incubated overnight at 4°C in the primary antibody. The following primary antibodies were used: rabbit MECP2 (Diagenode C15410052, 1:1000), rabbit OCT1 (Novus Biologicals NBP2-21584, 1:500), rabbit pATR (Abcam ab227851 1:500). Next, the slides were washed three times with 0.1% PBST and secondary antibody conjugated with Alexa 488, 568, 594 or 647 (1:500, Life Technologies A-21203, A21202, A31571, A-21207) was used, accompanied by DAPI (Invitrogen 1:500). Slides were then washed three times with 0.1% PBST and mounted using Prolong Gold (Invitrogen). Mean fluorescent intensity was quantified using ImageJ.

### Cytospin

Sorted nuclei were injected into the cytofunnel and spun down at 600 rpm for 15 minutes. Next, the nuclei were fixed with 4% paraformaldehyde following the immunofluorescence procedure.

### UV irradiation

Cell culture media was exchanged for PBS and the cells were placed in the GS GENE LINKER UV chamber (Bio-Rad). The irradiation was set to 100 mJ for 45 seconds. Next, the cells were washed with PBS and growth factor withdrawal media was applied. 4 hours later the cells were fixed for immunofluorescence.

### Disruption of mitochondrial function

Cells were washed with PBS and treated with either DMSO, 1 uM Phenformin, 20 uM Phenformin, 0.1 uM Rotenone or 5 uM Rotenone for three days. Media was changed every day.

### C13 labeled glucose incorporation

Cells were fed with DMEM (ThermoFisher Scientific) supplemented with 4 mM glutamine (ThermoFisher Scientific), 1 mM pyruvate (ThermoFisher Scientific), and 10 mM C13 labeled glucose (Cambridge Isotope Laboratories). 24 hours later the cells were washed twice with ammonium acetate (ThermoFisher Scientific) on ice and 80% methanol (ThermoFisher Scientific) was added to the cells. The plates were placed in −80C for 15 minutes. Next, the cells were scraped off the plate into eppendorf tubes, vortexed and centrifuged at 17000 g for 10 minutes at 4C. The methanol was then evaporated using the EZ-Lite evaporator. Dried samples were analyzed using Mass Spectrometry.

### Seahorse Assay

Cell were plated at a density 50,000-90,000 cells per well in a XF96 microplate (Agilent) and placed in the 37oC 5% CO2 incubator overnight. The next day, the cells were washed twice with the assay medium (Dulbecco′s Modified Eagle′s Medium supplemented with 10 mM glucose, 2 mM L-glutamine, 1 mM pyruvate and 5 mM HEPES, pH 7.4), and the microplate was placed in a 37oC incubator without CO2. 30 minutes later, the plate was loaded into the Seahorse XF96 Extracellular Flux Analyzer (Agilent Technologies). The following compounds were Injected during the assay: 2 uM oligomycin, 0.75 and 1.35 uM FCCP; 2 uM rotenone and antimycin A. When the measurements were done, cells were fixed with 4% paraformaldehyde, stained with Hoechst, and cell number per well was determined using an Operetta High-Content Imaging System (PerkinElmer). Oxygen consumption rates (OCR) were normalized to cell number per well.

### Nuclei isolation from frozen brain samples

Nuclei were isolated as described previously (Krishnaswami et al. 2016). Briefly, brain tissue was cut on ice using a scalpel and homogenized in a glass dounce. The homogenate was then filtered through a 40 um cell strainer and centrifuged at 1000g for 8 min at 4°C. The pellet was resuspended in a homogenization buffer and mixed with equal amounts of iodixanol. This mixture was then gently placed in a new tube over a 29% iodixanol. The nuclei were centrifuged at 13500g for 20 min at 4°C. The pellet was resuspended in the immunostaining buffer and incubated for 15 min at 4°C. Next, primary antibody was added to the nuclei pellet and incubated on a rotator at 4°C for 40 min. The following primary antibodies were used: rabbit MECP2 (Diagenode C15410052, 1:250), chicken Neun (Millipore Sigma MAB377 1:250). Nuclei were centrifuged at 400 g for 5 min at 4 °C and washed with PBS/BSA twice. Secondary antibodies conjugated with Alexa 488 and 594 (1:500, Life Technologies A-21203, A21202) were added, accompanied by DAPI (Invitrogen 1:500), the nuclei were incubated on a rotator at 4°C for 30 min. Immunostained nuclei were subjected to FACS.

### Library Preparation and Sequencing

Sorted nuclei were delivered to the UCLA Technology Center for Genomics × Bioinformatics where libraries for RNA sequencing were prepared. The samples were sequenced using NovaSeq 6000 S2 PE 2×50 with 50,000 reads per cell.

### Single-cell RNA-sequencing analysis

Aligned counts were clustered using Seurat v3 and the standard clustering pipeline and parameters. Clustermarkers were calculated using the Wilcoxon rank sum test to compare distributions across individual clusters and the rest of the population. All differential expression analyses were also calculated using the Wilcoxon rank sum test. Venn diagrams were generated using bioVenn. Cite: BioVenn – a web application for the comparison and visualization of biological lists using area-proportional Venn diagrams T. Hulsen, J. de Vlieg and W. Alkema, BMC Genomics 2008, 9 (1): 488. Gene ontology visualizations were generated using ggplot2, with geometric points and a scaled color gradient.

### Transcription factor binding site enrichment analysis

Analysis was performed using the DIRE tool https://dire.dcode.org.

### Gene ontology analysis

Analysis was performed using Metascape, a Gene Annotation & Analysis Resource https://metascape.org/gp/index.html#/main/step1.

## Supplemental Figure Legends

**Supplemental Figure 1. Quality control analysis of the single cell data** (**A**) A violin plot showing the number of counts per cell in the Drop Seq and 10x data. (**B**) Number of genes per cell in each sample. (**C**) Percent mitochondrial reads per cell in each sample. (**D**) Number of cells detected in each sample.

**Supplemental Figure 2. Reanalysis of published sc-RNA-seq data** (**A**) Left, the diagram depicts the process of nuclei isolation employed by the Greenberg group followed by immunostaining and FACS. Right, UMAP showing 12 unique clusters identified during the reanalysis of the data. (**B**) GO analysis of misregulated pathways shows similar trend to our 10x data, with synaptic and metabolic gene expression being altered due to lack of MECP2. (**C**) GO analysis of DEGs from mouse knock out model shows downregulation of genes related to synapse while inflammation related genes are upregulated due to lack of Mecp2.

**Supplemental Figure 3. POU2F1/OCT1 is one of the most enriched potential regulators of genes altered in the Aging brain** DIRE analysis on RNA-seq data from the Aging brain showed enrichment for OCT1 binding sites in DEGs.

