## Supplementary figures and images for "Defining dysfunction due to loss of MECP2 in Rett Patient Brain"

### Supplemental Figure 1

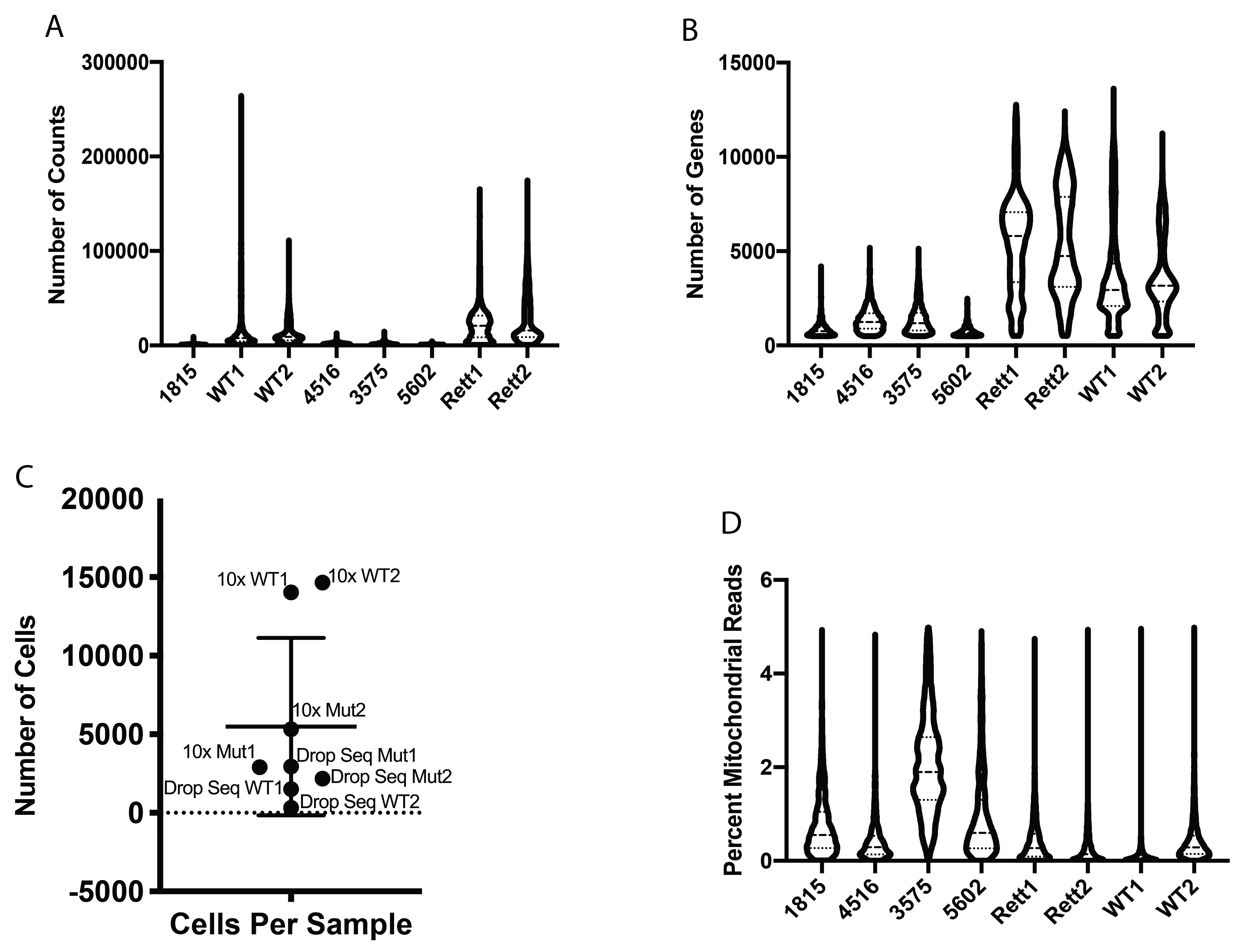

### Supplemental Figure 2

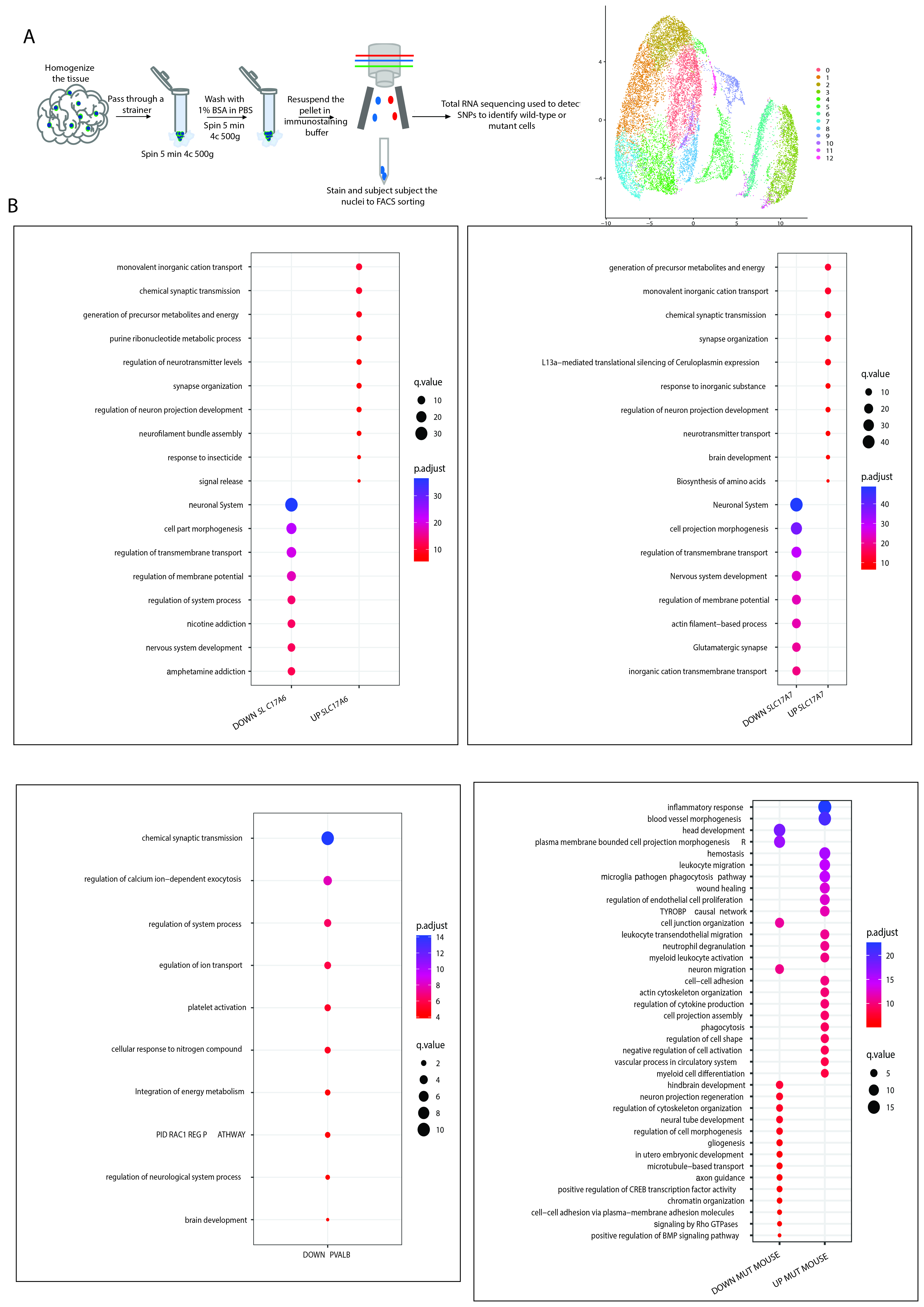

### Supplemental Figure 3

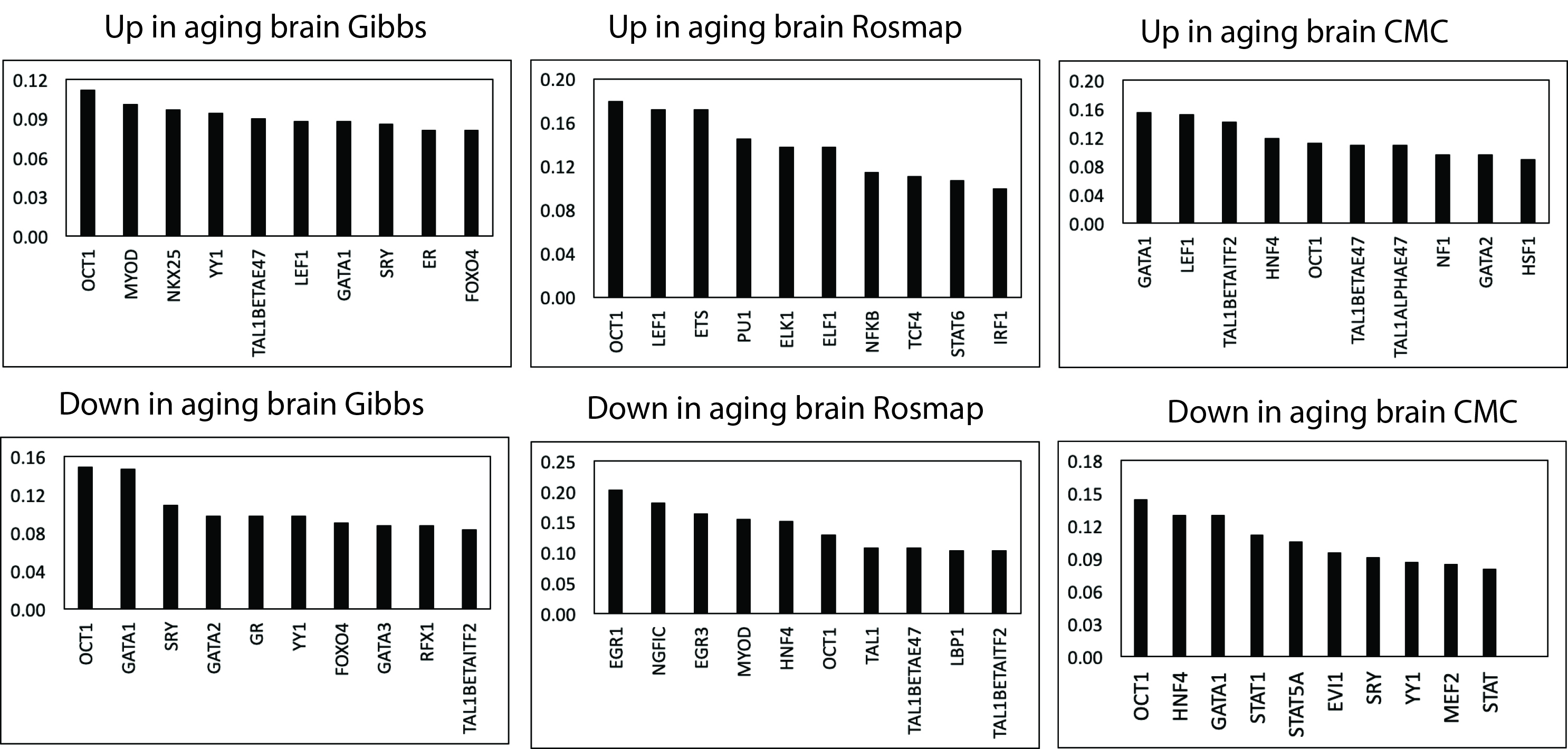
